# Prophylactic vaccination protects against the development of oxycodone self-administration

**DOI:** 10.1101/210369

**Authors:** Jacques D. Nguyen, Candy S. Hwang, Yanabel Grant, Kim D. Janda, Michael A. Taffe

**Affiliations:** Department of Neuroscience; Departments of Chemistry and Immunology, The Skaggs Institute for Chemical Biology, Worm Institute for Research and Medicine (WIRM); The Scripps Research Institute; La Jolla, CA, USA

**Author notes:** these authors contributed equally (Jacques D. Nguyen, Candy S. Hwang). Address Correspondence to: Dr. Michael A. Taffe, Department of Neuroscience, SP30–2400; 10550 North Torrey Pines Road; The Scripps Research Institute, La Jolla, CA 92037; USA.

**Keywords:** Oxycodone, vaccine, self-administration

## Abstract

Abuse of prescription opioids is a growing public health crisis in the United States, with drug overdose deaths increasing dramatically over the past 15 years. Few preclinical studies exist on the reinforcing effects of oxycodone or on the development of therapies for oxycodone abuse. This study was conducted to determine if immunopharmacotherapy directed against oxycodone would be capable of altering oxycodone-induced antinociception and intravenous self-administration. Male Wistar rats were administered a small-molecule immunoconjugate vaccine (Oxy-TT) or the control carrier protein, tetanus toxoid (TT), and trained to intravenously self-administer oxycodone (0.06 or 0.15 mg/kg/infusion). Brain oxycodone concentrations were 50% lower in Oxy-TT rats compared to TT rats 30 minutes after injection (1 mg/kg, s.c.) whereas plasma oxycodone was 15-fold higher from drug sequestration by circulating antibodies. Oxy-TT rats were also less sensitive to 1–2 mg/kg, s.c. oxycodone on a hot water nociception assay. Half of the Oxy-TT rats failed to acquire intravenous self-administration under the 0.06 mg/kg/infusion training dose. Oxycodone self-administration of Oxy-TT rats trained on 0.15 mg/kg/infusion was higher than controls; however under progressive ratio (PR) conditions the Oxy-TT rats decreased their oxycodone intake, unlike TT controls. These data demonstrate that active vaccination provides protection against the reinforcing effects of oxycodone. Anti-oxycodone vaccines may entirely prevent repeated use in some individuals who otherwise would become addicted. Vaccination may also reduce dependence in those who become addicted and therefore facilitate the effects of other therapeutic interventions which either increase the difficulty of drug use or incentivize other behaviors.

## 1. Introduction

Drug overdose deaths in the US have increased dramatically over the past 15 years with most of these attributable to opioids, including prescription drugs. Despite an increase in nonmedical prescription opioid abuse, with the use of heroin and other illicit drugs most often presaged by prescription opioids in new users (Dertadian and Maher, 2014; Mars *et al*, 2014), preclinical research into the addictive properties of oxycodone has been infrequent in comparison with studies on heroin or morphine. Limited studies in mice (Enga *et al*, 2016; Mayer-Blackwell *et al*, 2014; Zhang *et al*, 2015; Zhang *et al*, 2009) and rats (Leri and Burns, 2005; Mavrikaki *et al*, 2017; Secci *et al*, 2016; Wade *et al*, 2015) affirm that oxycodone will reinforce operant responding in intravenous self-administration (IVSA) models, thus, traditional pre-clinical measures of opioid addiction are useful to study the effects of potential therapeutics.

Anti-drug vaccination alters the effects of many drugs of abuse (Bremer and Janda, 2017; Lockner and Janda, 2013; Ohia-Nwoko *et al*, 2016); vaccines against cocaine (Haney *et al*, 2010; Kosten *et al*, 2002; Martell *et al*, 2009) and nicotine (Cornuz *et al*, 2008; Hatsukami *et al*, 2011) have advanced to clinical trials. Initial studies by Pravetoni and colleagues showed that an oxycodone-specific vaccine could induce antibodies capable of sequestering oxycodone in the blood and lowering brain penetration of oxycodone in rats (Pravetoni *et al*, 2012; Pravetoni *et al*, 2014). Vaccination decreased the antinociceptive effects of a single dose of oxycodone and reduced oxycodone intake during the acquisition of IVSA. Vaccination has also been shown to prevent oxycodone-induced respiratory depression and cardiovascular effects in rats (Raleigh *et al*, 2018; Raleigh *et al*, 2017). This success encourages more comprehensive research on anti-oxycodone vaccines, to further general understanding of how anti-drug vaccinations function in addition to supporting an eventual clinical vaccine for oxycodone abuse.

Janda and colleagues created a vaccine incorporating an oxycodone-hapten conjugated to tetanus toxoid (Oxy-TT) that has shown efficacy in attenuating antinociceptive effects and oxycodone overdose in mice (Kimishima *et al*, 2017). Therefore, it is of interest to determine if the Oxy-TT can reduce voluntary drug-seeking in a rat model. Prior evaluation of anti-drug vaccination efficacy in drug self-administration has typically focused on the effects of vaccination after the acquisition of drug taking behavior, e.g., for vaccines directed against cocaine (Carrera *et al*, 2000; Evans *et al*, 2016; Kantak *et al*, 2000), nicotine (Lindblom *et al*, 2002; Moreno *et al*, 2010) and morphine or heroin (Anton and Leff, 2006; Hwang *et al*, 2018a; Hwang *et al*, 2018b; Schlosburg *et al*, 2013). Although there have been a few studies of the effects of prophylactic vaccination prior to self-administration of cocaine (Wee *et al*, 2012), nicotine (LeSage *et al*, 2006), oxycodone (Pravetoni *et al*, 2014), heroin (Stowe *et al*, 2011) and methamphetamine (Duryee *et al*, 2009; Miller *et al*, 2015), the effects of vaccination in the early stages of addiction are not well understood. Prior studies of IVSA of oxycodone (Pravetoni *et al*, 2014) and methamphetamine (Duryee *et al*, 2009) suggest that increasing the workload (i.e., the number of lever responses required for each infusion) may decrease intake in the vaccinated groups relative to controls. Another finding with an anti-methamphetamine vaccine suggests, however, that initial higher drug intake may be gradually extinguished over the first several IVSA sessions with no change in workload (Miller *et al*, 2015). Cocaine IVSA is lower in vaccinated animals than in controls under a Progressive Ratio (PR) schedule of reinforcement and increased workload (Wee *et al*, 2012), whereas nicotine IVSA *increases* in the vaccinated rats (LeSage *et al*, 2006). The present study was conducted to determine if the novel Oxy-TT vaccine directed against oxycodone would be capable of altering the IVSA of oxycodone.

## 2. Materials and Methods

### 2.1 Animals

Adult male Wistar (N=48; Charles River, New York) rats were housed in humidity and temperature-controlled vivaria (23±1^°^C) on 12:12 hour light:dark cycles. Cohort 1 (N=24; lower training dose) animals were 10 weeks old (mean weight 288.3 g; SD 12.8) at the start of the study. Cohort 2 (N=24; higher training dose) animals were 13 weeks old (mean weight 408.9 g; SD 30.6) at the start of the study. Animals had *ad libitum* access to food and water in their home cages. All procedures were conducted under protocols approved by the Institutional Animal Care and Use Committees of The Scripps Research Institute and in a manner consistent with the National Institutes of Health Guide for the Care and Use of Laboratory Animals.

### 2.2 Hapten Synthesis

The oxycodone hapten (Oxy) was designed with an activated linker extending from the bridgehead nitrogen to directly react with the surface lysines of carrier protein tetanus toxoid (TT, **Figure 1A**) or BSA. Compound **7** was synthesized according to previously published methods in our laboratory with slight modification in the reductive amination and amide bond formation steps (Kimishima *et al*, 2016). Although decreasing the equivalents of NaBH(OAc)_3_ to 1.1 equivalents avoids formation of the undesired alcohol, the reaction results in an incomplete mixture of **2** and **4** after several hours. Instead, 3 equivalents of NaBH(OAc)_3_ rapidly led to the conversion of the reductive amination and reduced C-6 ketone product. The secondary C-6 alcohol can be simply and mildly oxidized to the desired ketone (**4**) using DMP.(Kimishima *et al*, 2014) After preparation and confirmation of the product (**4**) by ^1^H and ^13^C NMR, **4** was deprotected and condensed with **5** using HATU as a catalyst, resulting in a 61% yield of purified **6**. Additional details can be found in the Supplementary Information.

#### 2.2.1 Synthesis of tert-butyl(4-((4R,4aS,7aR,12bS)-4a-hydroxy-9-methoxy-7-oxo-1,2,4,4a,5,6,7,7a-octahydro-3H-4,12-methanobenzofuro[3,2-e]isoquinolin-3-yl)butyl)carbamate (**4**)

Oxycodone hydrochloride (100 mg, 0.28 mmol) was dissolved in dichloromethane (CH_2_Cl_2_) and washed with saturated sodium bicarbonate (2 × 10 mL). The solvents from the organic layer were dried with sodium sulfate (Na_2_SO_4_), filtered and evaporated. The white powder was dissolved in 4 mL of dry, 1,2-dichloroethane. Sodium bicarbonate (235 mg, 2.8 mmol, 10 equiv) and ACE-Cl (245 µL, 2.3 mmol, 8 equiv) were added at room temperature (rt). The mixture was then heated to reflux overnight under argon and checked by TLC (5:1 CH_2_Cl_2_:methanol). The reaction was then cooled and solvent was removed under reduced pressure. The residue was then dissolved in 10 mL CH_2_Cl_2_ and washed with saturated bicarbonate (2 × 10 mL). The aqueous layers were combined and washed with ethyl acetate (EtOAc, 1 × 10 mL). The organic layers were combined and dried with Na_2_SO_4_. The solution was filtered and solvents were evaporated. The oil was then dissolved in a portion of methanol (MeOH), stirred at rt for several hours and monitored by TLC (5:1 CH_2_Cl_2_:MeOH). The solvents were evaporated and the residue was purified by flash chromatography using 10% MeOH in CH_2_Cl_2_ to give 36 mg of **2** (43% crude yield) as a colorless oil. ESI-MS: MS (*m*/*z*): *calcd* for C_17_H_20_NO_4_^+^: 302.1, *found:* 302.2 [M + H]^+^.

Noroxycodone **2** (36 mg, 0.12 mmol) was dissolved in 4 mL of dry CH_2_Cl_2_. Na_2_SO_4_ (26 mg, 0.18 mmol, 1.5 equiv) was added, followed by dropwise addition of **3** (45 mg, 0.24 mmol, 2 equiv). The mixture was allowed to stir for twenty minutes before addition of NaBH(OAc)_3_ (76 mg, 0.36 mmol, 3 equiv) to initiate the reduction. The reaction was allowed to stir for 2 h and monitored by TLC (95:5 EtOAc:MeOH). After completion of the reaction, NaBH(OAc)_3_ was quenched with water and the reaction was washed with saturated sodium bicarbonate (2 × 10 mL). The organic layers were dried with Na_2_SO_4_ and solvents were evaporated. The residue was purified by flash chromatography using 5% MeOH in EtOAc. The fractions were collected and the solvents were evaporated to give 22 mg of the putative alcohol from ketone reduction at the C-6 position (39% yield), as confirmed by MS. ESI-MS: MS (*m/z*)*: calcd* for C_26_H_39_N_2_O_6_^+^: 475.3, *found:* 475.3 [M + H]^+^.

The alcohol (22 mg, 0.05 mmol) was dissolved in 3 mL of dry CH_2_Cl_2_ and sodium bicarbonate (8.4 mg, 0.10 mmol, 2 equiv) and DMP (30 mg, 0.07 mmol, 1.3 equiv) were added to initiate the reaction. The reaction was allowed to stir for several hours and monitored by TLC (9:1 CH_2_Cl_2_:MeOH) and LCMS. The mixture was diluted with CH_2_Cl_2_ and washed with 20 mL of a 1:1 solution of saturated bicarbonate and saturated sodium thiosulfate. The aqueous layer was washed with portions of a 9:1 CH_2_Cl_2_:MeOH solution (3 × 10 mL). The organic layers were collected and the solvents were evaporated. The residue was then dissolved in 10 mL of 9:1 CH_2_Cl_2_:MeOH and filtered through a short pad of silica. The eluted solvents were dried with Na_2_SO_4_ and the solvents were removed to give 20 mg of **4** (92% yield). NMR spectra were consistent with literature values.(Kimishima *et al*, 2016) ^1^H NMR (500 MHz, CDCl_3_) δ 6.70 (d, *J* = 8.2 Hz, 1H), 6.62 (d, *J* = 8.2 Hz, 1H), 4.67 (br, s, 1H), 4.61 (s, 1H), 3.90 (s, 3H), 3.35 (s, 1H), 3.19 − 3.12 (m, 2H), 3.10 (d, *J* = 18.8 Hz, 1H), 3.00 (td, *J* = 14.5, 5.2 Hz, 2H), 2.62 (dd, *J* = 18.3, 5.5 Hz, 2H), 2.58 − 2.53 (m, 2H), 2.44 (dd, *J* = 11.8, 4.2 Hz, 1H), 2.29 (d, *J* = 14.3 Hz, 1H), 2.18 (dt, *J* = 14.5, 7.1 Hz, 1H), 2.12 − 2.00 (m, 1H), 1.88 (d, *J* = 13.4 Hz, 1H), 1.70 − 1.51 (m, 3H), 1.45 (s, 9H). ^13^C NMR (126 MHz, CDCl_3_) δ 208.50, 145.18, 143.19, 129.46, 124.73, 119.59, 115.18, 90.44, 70.43, 62.99, 57.00, 54.04, 50.79, 43.81, 40.38, 36.21, 31.63, 30.61, 28.58 (3C), 27.98, 24.71, 23.13. ESI-MS: MS (*m/z*): *calcd* for C_26_H_37_N_2_O_6_^+^: 473.3, *found:* 473.3 [M + H]^+^.

**Figure 1.**
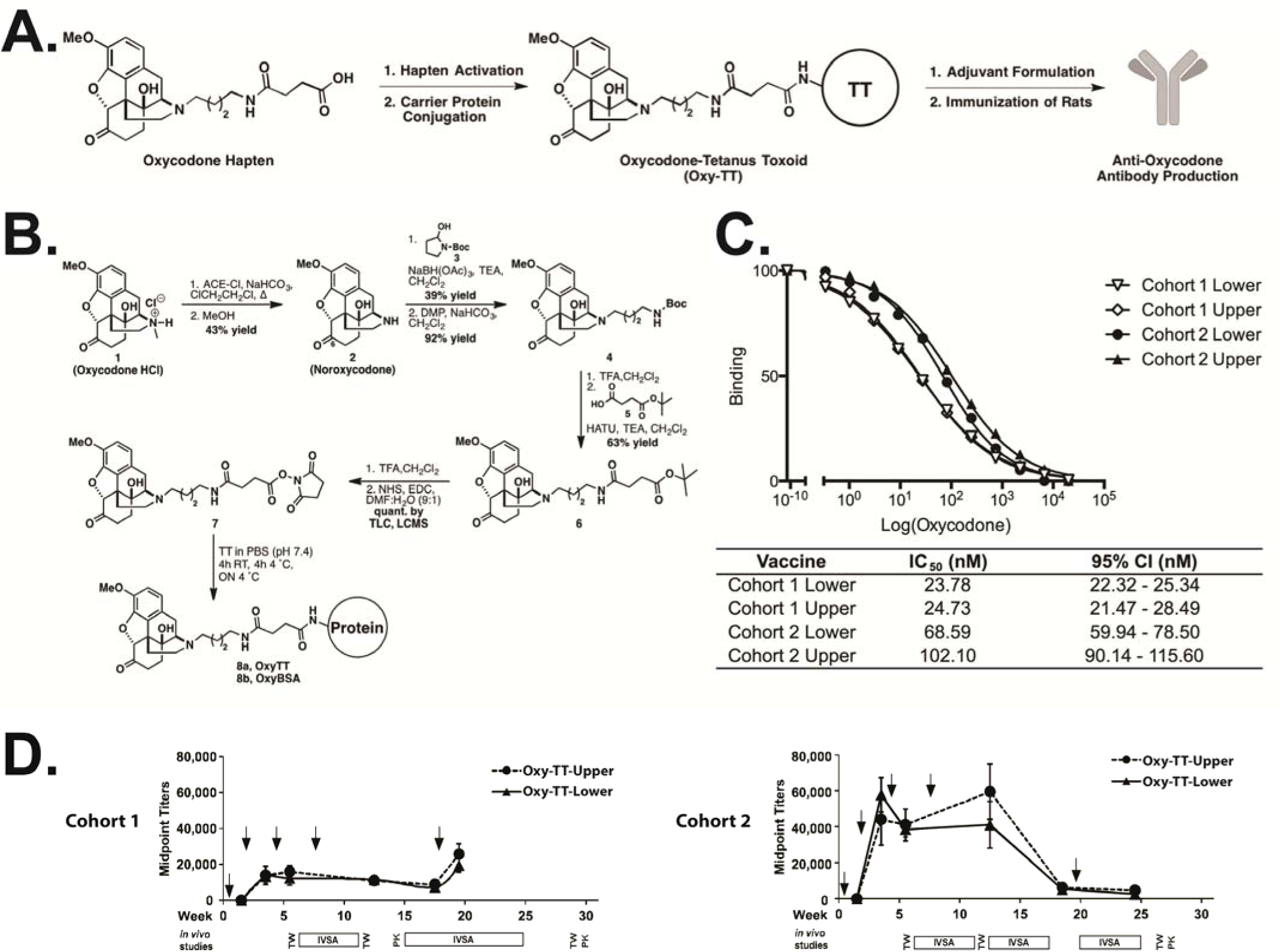
The general vaccination strategy using an oxycodone hapten-immunoconjugate vaccine. A) The general vaccine strategy and B) scheme of Oxy hapten synthesis and preparation of Oxy-TT and Oxy-BSA immunoconjugates. C) Surface plasmon resonance (SPR) analysis of binding interactions confirmed that the polyclonal response presented binding to free drug in Cohorts 1 and 2. D) The vaccination schedule resulted in plasma anti-Oxy antibody midpoint titers from Oxy-TT vaccinated rats. Cohort 1 and Cohort 2 are separated into Upper or Lower groups (N=5–6) with alum and CpG ODN 1826 as adjuvants as determined by ELISA. Arrows indicate when rats were vaccinated. Data are presented as means ± SEM. Plasma from control TT vaccinated rats with alum and CpG ODN 1826 as adjuvants did not contain any detectable anti-Oxy IgG titers. TW and PK indicated tail-withdrawal tests and blood-draws for pharmacokinetic analysis, respectively.

#### 2.2.2 Synthesis of tert-butyl 4-((4-((4R,4aS,7aR, 12bS)-4a-hydroxy-9-methoxy-7-oxo-1,2,4,4a,5,6,7,7a-octahydro-3H-4,12-methanobenzofuro[3,2-e]isoquinolin-3-yl)butyl)amino)-4-oxobutanoate **(6)**

Compound **4** (20 mg, 0.04 mmol) was deprotected using 2 mL of a 1:1 solution of TFA and CH_2_Cl_2_. The deprotection was allowed to stir for 2 h, and was monitored by TLC (9:1 CH_2_Cl_2_:MeOH). After complete deprotection of the Boc group, the solvents were coevaporated with several portions of toluene and CH_2_Cl_2_. The compound was then dissolved in 1 mL of dry CH_2_Cl_2_ and 18 µL of TEA (0.13 mmol, 13 mg, 3.3 equiv). Mono-t-butyl succinate **5** (0.05 mmol, 9 mg, 1.1 equiv) and HATU (0.05 mmol, 19 mg, 1.1 equiv) were added in one portion to the solution. The reaction was allowed to stir for 3 h and monitored by TLC (9:1 CH_2_Cl_2_:MeOH). After complete formation of the amide (**6**), the reaction was diluted with 10 mL of CH_2_Cl_2_ and washed with saturated sodium bicarbonate. The organic layers were dried with Na_2_SO_4_, filtered and the solvents were evaporated. The crude oil was purified by flash chromatography using 10% MeOH in CH_2_Cl_2_. The pure fractions were combined and solvents were evaporated to yield 14 mg of **6** (63% yield). ^1^H NMR (500 MHz, CDCl_3_) δ 6.69 (d, *J* = 8.2 Hz, 1H), 6.61 (d, *J* = 8.2 Hz, 1H), 4.65 (s, 1H), 3.89 (s, 3H), 3.27 (q, *J* = 6.5 Hz, 2H), 3.08 (d, *J* = 18.5 Hz, 1H), 3.01 (td, *J* = 14.4, 5.0 Hz, 1H), 2.96 (d, *J* = 5.9 Hz, 1H), 2.64 – 2.46 (m, 7H), 2.41 (t, *J* = 6.8 Hz, 2H), 2.37 (d, *J* =7.4 Hz, 1H), 2.28 (dt, *J* = 14.4, 3.2 Hz, 1H), 2.14 (td, *J* = 12.1, 3.7 Hz, 1H), 1.86 (ddd, *J* = 13.4, 5.0, 2.9 Hz, 1H), 1.68 – 1.56 (m, 3H), 1.43 (s, 9H). ^13^C NMR (126 MHz, CDCl_3_) δ 208.60, 172.59, 171.96, 145.15, 143.12, 129.54, 124.92, 119.55, 115.09, 90.45, 80.95, 70.43, 63.09, 56.98, 53.93, 50.85, 43.62, 39.33, 36.25, 31.60, 31.52, 31.05, 30.73, 28.22 (3C), 27.51, 24.85, 23.09. ESI-MS: MS (*m/z*): *calcd* for C_29_H_41_N_2_O_7_^+^: 529.3, *found:* 529.3 [M + H]^+^.

#### 2.2.3 Synthesis of activated oxycodone hapten **7** and conjugation to carrier protein tetanus toxoid (TT) or bovine serum albumin (BSA)

To a 10 mg aliquot of **6** (0.019 mmol) was added 1 mL of a mixture of TFA and CH_2_Cl_2_ (3:1). The deprotection was allowed to proceed for several hours and was monitored by TLC (9:1 CH_2_Cl_2_:MeOH) and LCMS. ESI-MS: MS (*m/z*): *calcd* for C_25_H_33_N_2_O_7_^+^: 473.2, *found:* 473.2 [M + H]^+^. The deprotected hapten was then coevaporated with several portions of toluene and CH_2_Cl_2_. The deprotected acid was dissolved in 1000 µL of a 9:1 DMF:H_2_O solution, followed by addition of TEA (0.06 mmol, 7.9 µL, 3 equiv). NHS (0.19 mmol, 22 mg, 10 equiv) and *N*-(3-dimethylaminopropyl)-*N*’-ethylcarbodiimide hydrochloride (EDC, 0.09 mmol, 17.3 mg, 10 equiv) were added in one portion. The solution was allowed to stir for an hour and was monitored by LCMS. Another 5 equiv of NHS and EDC were added in one portion to the reaction. After one hour, LCMS revealed completion of the reaction. ESI-MS: MS (m/z): *calcd* for C_32_H_38_N_3_O_10_^+^: 570.2, *found:* 570.3 [M + H]^+^.

### 2.3 MALDI-ToF Analysis

Oxycodone immunoconjugated to BSA (OxyBSA) was used as a surrogate for TT to quantify the number of oxycodone haptens (Oxy) and for ELISAs. The immunoconjugate was run through a PD MiniTrap G-10 desalting column (GE Healthcare) and then analyzed by MALDI-ToF for the hapten:carrier protein conjugation number. In order to quantify the copy number or the number of oxycodone haptens (Oxy) on BSA, the molecular weight of conjugated BSA (OxyBSA, MW: 73,890 by MALDI-ToF) was compared to the molecular weight of unconjugated BSA (BSA, MW: 66,347 by MALDI-ToF). Additional details can be found in the Supplemental Information.

### 2.4 Vaccine Formulation

Vaccines were formulated the day of immunization using 13:1:5 (v/v/v) mixture of Oxy-TT (1.0 mg/ml in PBS) or control TT (1.0 mg/ml in PBS), CpG ODN 1826 (5 mg/ml in PBS), and Alhydrogel^®^ (alum, 10 mg/ml, InvivoGen). Rats were administered the conjugate vaccine (Oxy-TT; N=24) or tetanus toxoid only control (TT; N=24) on Weeks 0, 2, 4, 8 and 18 for Cohort 1 and on Weeks 0, 2, 4, 8 and 20 for Cohort 2 (**Figure 1D**). The immunization protocol was adapted from a vaccination protocol previously reported (Nguyen *et al*, 2016; Nguyen *et al*, 2017a). Vaccines were formulated the day of immunization using 13:1:5 (v/v/v) mixture of Oxy-TT (1.0 mg/ml in PBS) or control TT (1.0 mg/ml in PBS), CpG ODN 1826 (5 mg/ml in PBS), and Alhydrogel^®^ (alum, 10 mg/ml, InvivoGen). CpG ODN 1826 is a phosphorothionated oligonucleotide and murine TLR9 agonist with the following sequence (5’ to 3’): TCCATGACGTTCCTGACGTT (Eurofins MWG Operon) (Bremer *et al*, 2014). Each vaccine was prepared by shaking the mixture for thirty minutes prior to injection. The delivered dose of each component was 200 μg Oxy-TT or TT, 150 μg of CpG ODN 1826, and 1.5 mg of alum per animal for each intraperitoneal injection (i.p.). Additional details can be found in the Supplemental Information.

### 2.5 Plasma Titer Analysis

Rats were anesthetized with an isoflurane/oxygen vapor mixture (isoflurane 5% induction, 1–3% maintenance), and blood was collected from the jugular vein on Weeks 3, 5, 12, 17 and 19 and for Cohort 1 and on Weeks 3, 5, 12, 18 and 24 for Cohort 2. Titers were analyzed by ELISA and defined by the dilution required to achieve a 50% signal (midpoint titer). Absorbance values were normalized to the highest absorbance value per set, and a curve was fit using the log(inhibitor) vs. Normalized response – variable slope equation to determine midpoint titer and standard error (GraphPad PRISM). Plasma from control TT rats did not contain any detectable anti-oxycodone titers. Data were tested for statistical outlier evaluation using Grubbs test and no outliers were detected.

Costar 3690 plates with half-area, high-binding 96-well microtiter plates were coated with 50 ng of OxyBSA (25 µL of 2 µg/mL OxyBSA conjugate in PBS) per well overnight at 37 °C. The plates were blocked with 5% skim milk in PBS buffer (pH 7.4) for one hour at rt. Vaccinated rat plasma was serially diluted 1:1 in 2% BSA in PBS buffer (pH 7.4) across twelve columns starting at 1:200. After two hours of incubation at room temperature, the plates were washed ten times with PBS, and goat anti-rat IgG horseradish peroxidase (HRP) secondary antibody (Southern Biotech) at 1:10,000 dilution in 2% BSA in PBS buffer (pH 7.4) was added and incubated for two hours at room temperature. After washing ten times with PBS buffer, a 1:1 solution of 3,3’,5,5’-tetramethylbenzidine (TMB) and H_2_O_2_ substrate (Thermo Pierce) were added. Plates were incubated for 15 minutes and then quenched with 2.0 M H_2_SO_4_. Plates were read at 450 nm. Details of determination of binding affinity of antibodies by surface plasmon resonance can be found in the Supplemental Information.

### 2.6 Drugs

Oxycodone hydrochloride used for this study was obtained from Sigma-Aldrich (St. Louis, MO) and 3,6-Diacetylmorphine HCl (Heroin HCl) was obtained from NIDA Drug Supply. Drugs were expressed as the salt and dissolved in physiological saline (0.9%). For tail-withdrawal studies, rats were administered 0.9% saline vehicle, oxycodone (0.5, 1.0, 2.0 mg/kg) or heroin (1.0 mg/kg) via subcutaneous (s.c.) injection.

### 2.7 Intravenous Catheterization and Self-Administration

Rats were anesthetized with an isoflurane/oxygen vapor mixture (isoflurane 5% induction, 1–3% maintenance) and prepared with chronic intravenous catheters as described (Miller *et al*, 2013; Nguyen *et al*, 2017b) on week 4 of the vaccination protocol. The catheters consisted of a 14.5-cm length polyurethane based tubing (MicroRenathane^®^, Braintree Scientific, Inc, Braintree MA, USA) fitted to a guide cannula (Plastics one, Roanoke, VA) curved at an angle and encased in dental cement anchored to an ~3-cm circle of durable mesh. Catheter tubing was passed subcutaneously from the animal’s back to the right jugular vein. Catheter tubing was inserted into the vein and secured gently with suture thread. A liquid tissue adhesive was used to close the incisions (3M™ Vetbond™ Tissue Adhesive; 1469S B). A minimum of 4 days was allowed for surgical recovery prior to starting an experiment. For the first three days of the recovery period, an antibiotic (cephazolin) and an analgesic (flunixin) were administered daily. During testing and training, intravenous catheters were flushed with ~0.2–0.3 ml heparinized (32.3 USP/ml) saline before sessions and ~0.2–0.3 ml heparinized saline containing cefazolin (100 mg/ml) after sessions. Catheter patency was assessed weekly after the first two weeks via administration through the catheter of ~0.2 ml (10 mg/ml) of the ultra-short-acting barbiturate anesthetic, Brevital sodium (1% methohexital sodium; Eli Lilly, Indianapolis, IN). Animals with patent catheters exhibit prominent signs of anesthesia (pronounced loss of muscle tone) within 3 s after infusion. Animals that failed to display these signs were considered to have faulty catheters and were discontinued from the study. Data that was taken prior to failing this test and after the previous passing of this test were excluded from analysis.

Drug self-administration was conducted in operant boxes (Med Associates) located inside sound-attenuating chambers located in an experimental room (ambient temperature 22±1^°^C; illuminated by red light) outside of the housing vivarium. To begin a session, the catheter fittings on the animals’ backs were connected to polyethylene tubing contained inside a protective spring suspended into the operant chamber from a liquid swivel attached to a balance arm. Each operant session started with the extension of two retractable levers into the chamber. Following each completion of the response requirement (response ratio), a white stimulus light (located above the reinforced lever) signaled delivery of the reinforcer and remained on during a 20 s post-infusion timeout, during which responses were recorded but had no scheduled consequences. Drug infusions were delivered via syringe pump. The training doses (Cohort 1: 0.06 mg/kg/infusion; Cohort 2: 0.15 mg/kg/infusion) and session duration of 1 h were selected from a prior self-administration study (Wade et al. 2014). Rats were allowed to self-administer under a Fixed Ratio 1 (FR1) contingency for 18 sessions. Successful acquisition in the first 18 sessions was defined as an average of 7 or more infusions across two sequential days; the day of acquisition was defined as the first day. Significant individual differences were shown to contribute to outcome in human clinical trials of cocaine and nicotine vaccines (Hoogsteder *et al*, 2014; Martell *et al*, 2009). Likewise, there are significant individual differences in drug use trajectories, since problematic drug use is a minority outcome among those who sample drugs (Anthony *et al*, 1994; Schramm-Sapyta *et al*, 2009; Taffe, 2015). thus a median split analysis was planned a priori to address the possibility of individual differences of outcome in this study. Median splits were determined by ranking animals on their mean number of infusions obtained across 18 sessions, and are referred to as Upper and Lower. The 19^th^ session was a PR session with the respective training dose for all animals. In the PR paradigm, the required response ratio is increased after each reinforcer delivery within a session (Hodos, 1961; Segal and Mandell, 1974) as determined by the following equation (rounded to the nearest integer): Response Ratio=5e^^^(injection number\**j*)–5 (Richardson and Roberts, 1996). The *j* value is as specified for a given experiment, i.e., PR^J2^ refers to a session with the *j* value set to 0.2.

<u>Cohort 1</u>: Sessions 20–21 were FR1/training dose and Session 22 featured saline substitution. Training conditions were restored for Sessions 28–30 (i.e., FR1) and thereafter the schedule was FR5 for Sessions 31–44 and FR10 for Sessions 45–63. The PR^J2^ schedule was in effect for Sessions 64–70 and FR1 for Sessions 71–73. Mean infusions in Sessions 28–30 were used to re-rank individuals for the median split for this part of the experiment to determine effects of schedule changes on current, rather than acquisition, drug preference phenotype. This resulted in 3 individuals in the original lower half of the TT group (TT Lower) switching with Upper half individuals and one individual in the original Oxy-TT Lower half switching with an Upper half individual.

<u>Cohort 2</u>: Sessions 20–23 were PR^J2^ sessions with doses (0.0, 0.06, 0.15, 0.3 mg/kg/inf) presented in randomized order. There was negligible difference attributable to dose and the subsequent experiments focused on FR/PR transitions at a fixed per-infusion dose. Sessions 24–25 restored the training dose under FR1. Sessions 26–30 were PR^J2^ and sessions 31–35 were PR^J3^ (0.15 mg/kg/inf). Sessions 36–37 were FR1 (0.15 mg/kg/inf). Sessions 38–39 were FR1 (0.06 mg/kg/inf) and session 40 was PR^J3^ (0.06 mg/kg/inf).

### 2.8 Nociception

Tail withdrawal latency was assessed before, and 30 minutes after, injection of oxycodone (0.5, 1.0 or 2.0 mg/kg, s.c.) or heroin (1.0 mg/kg, s.c.) using a 52^°^C water bath and a 15 s maximum interval (Gold et al., 1994). Studies were conducted across the course of IVSA (pre- and postacquisition training) to verify *in vivo* efficacy of the vaccine. Tests were performed on Weeks 5, 10 and 27 (data not included) for Cohort 1 and on Weeks 5, 9, 11 (heroin/oxycodone) and 25 (data not included) for Cohort 2.

### 2.9 Determination of Plasma and Brain Oxycodone

In Cohort 1, Oxy-TT (N=11) and TT rats (N=11) blood was collected 30 minutes after 1.0 mg/kg oxycodone, s.c., on Week 14 (6 weeks from last boost). Blood and brain tissue were collected 30 minutes after 1.0 mg/kg oxycodone, s.c., (N=5, Oxy-TT; N=6, TT) or 2.0 mg/kg oxycodone, s.c., (N=5 per group) on Week 29. In Cohort 2, blood was collected from rats (N=12, Oxy-TT; N=10, TT) 30 minutes after 2.0 mg/kg oxycodone, s.c. on Week 26 (6 weeks from last boost). In Week 27, blood was collected 15 and 30 minutes after 1.0 mg/kg oxycodone, s.c., and brains were collected at the 30 minute time point. Two rats from the TT control group were not given oxycodone in order to use their blood and brain tissue as blank controls for generating the calibration curves. One rat from the Oxy-TT group was excluded from a 15-minute time point for the 1.0 mg/kg dose due to timing, but blood and brain tissue were harvested at 30 minutes for the same rat.

The rats were anesthetized and then rapidly decapitated using a sharp guillotine. The brain and trunk blood were collected. The blood was collected in tubes coated with lithium heparin and placed on ice, centrifuged at 10,000 rpm for 10 minutes, and then the plasma was collected. Brain tissue was immediately flash frozen using a dry ice/isopentane bath. Plasma and brain tissue were stored at −80 °C. Each frozen brain was weighed and an equivalent volume of 0.2 M aqueous perchloric acid (1 mL per gram) was added. The brain tissue was homogenized using a Bullet Blender with zirconium oxide beads (0.5 mm diameter, Thomas Scientific) and then centrifuged at 3,000 rpm for ten minutes. A 200 µL aliquot of the supernatant was added to 100 µL of spiked oxycodone concentrations (for standard curve, made up in H_2_O) or 100 µL of H_2_O (for samples) and 40 µL of *d*_6_-oxycodone (*d*_6_-Oxy, 1 µg/mL in MeOH). The mixture was vortexed for a minute to equilibrate, and then extracted with the Oasis PRiME HLB Extraction Cartridge. The eluted sample was evaporated using GENEVAC and taken up in MeOH for LCMS analysis.

To each 50 µL aliquot of plasma, 0.1 M perchloric acid (100 µL) was added, along with 20 µL of *d*_6_-oxycodone (1 µg/mL in MeOH) and 20 µL of spiked oxycodone concentrations or H_2_O. The mixture was vortexed for one minute and then run through the Oasis extraction plate. The eluent was evaporated and analyzed by LCMS. A standard curve was constructed using the ratio of peak area of drug analyte (Oxy) to the internal standard (*d*_6_-Oxy) versus concentration for both the brain and plasma samples. Linearity of the calibration curves were indicated by a correlation coefficient greater than 0.990 (R^2^ = 0.996 > 0.990; R^2^ = 0.993 > 0.990, respectively). Plasma and brain oxycodone levels were tested for statistical outlier evaluation using Grubbs test and significant outliers were removed. In the blood-brain distribution, only one outlier was detected from a brain sample from the Oxy-TT group (Z_crit_=2.411<2.58, N=12, *p*<0.05).

### 2.10 Data Analysis

Intravenous self-administration data were analyzed with repeated-measure Analysis of Variance (rmANOVA) with Vaccine Group (TT Upper/Lower and Oxy-TT Upper/Lower) as a between-subjects factor and Time (Session) as a within-subjects factor. Tail-withdrawal latencies were analyzed with rmANOVA with Vaccine as a between-subjects factor and with Time (and therefore Drug treatment) as a within-subjects factor. Plasma data were analyzed with two-way rmANOVA with vaccine Group as a between-subjects factor and Drug condition as the within-subjects factor as repeated measure factors. Brain concentrations were analyzed by unpaired t-tests. Significant effects within group were followed with post hoc analysis using Tukey correction for all multi-level factors and Sidak for two-level factors. Criterion for significant results was set at *p*<0.05, and all analysis used Prism for Windows (v. 6.07 and 7.03; GraphPad Software, Inc, San Diego CA).

## 3. Results

### 3.1 Conjugation of Oxy-TT Vaccine

The oxycodone hapten (Oxy) and Oxy-TT immunoconjugate (**Figure 1A**) were prepared as described previously (Kimishima *et al*, 2016) with minor modification detailed in the Experimental Section (**Figure 1B**). MALDI-ToF analysis of Oxy-BSA confirmed successful conjugation with approximately 16 moles of Oxy hapten per mole of carrier protein. Following the vaccination regimen, rats in the Oxy-TT groups established plasma antibody titer (**Figure 1D**). No anti-oxycodone Abs were detected in the control TT group.

### 3.2 Vaccination Protects Against Self-Administration of Oxycodone

Using a threshold per infusion dose of oxycodone (0.06 mg/kg/inf; Cohort 1) under fixed-ratio (FR) schedule of reinforcement, rats steadily increased intake across the first 18 sessions of training (**Figure 2A**; See details of FR schedule in Materials and Methods). Analysis of the data confirmed a significant effect of Session [F(17,357)=14.54; *p*<0.0001] but not of Group or of the interaction of factors. Only 58% of the Oxy-TT group met the acquisition criterion (average of 7 infusions across two sessions), whereas all of the TT animals met criterion (**Figure 2C**). Analysis of the acquisition data by median split (**Figure 2E**) confirmed a significant effect of Group [F(3,19)=7.83; *p*=0.0013], of Session [F(17,323)=16.95; p<0.0001] and of the interaction [F(51,323)=2.84; p<0.0001]. Collapsed across session, significantly more infusions were obtained by the Oxy-TT Upper group compared with each of the Lower half groups. Under a higher per infusion dose (0.15 mg/kg/inf; Cohort 2), rats increased oxycodone intake over the first 18 sessions of acquisition, with the Oxy-TT group eventually obtaining more infusions (**Figure 2B**). The analysis confirmed significant effects of Session [F(17,357)=19.89; *p*<0.0001], of vaccine Group [F(1,21)=6.02; *p*=0.0229] and of the interaction [F(17,357)=3.22; *p*<0.0001]. In this Cohort, 83% of the TT and Oxy-TT animals met acquisition criterion by averaging 7 infusions across two sessions (Figure 2D). Analysis of the acquisition data by median split (Figure 2F) confirmed a significant effect of Group [F(3,19)=11.72; *p*=0.0001], of Session [F(17,323)=19.05; *p*<0.0001] and of the interaction of factors [F(51,323)=1.62; *p*=0.0075].

**Figure 2.**
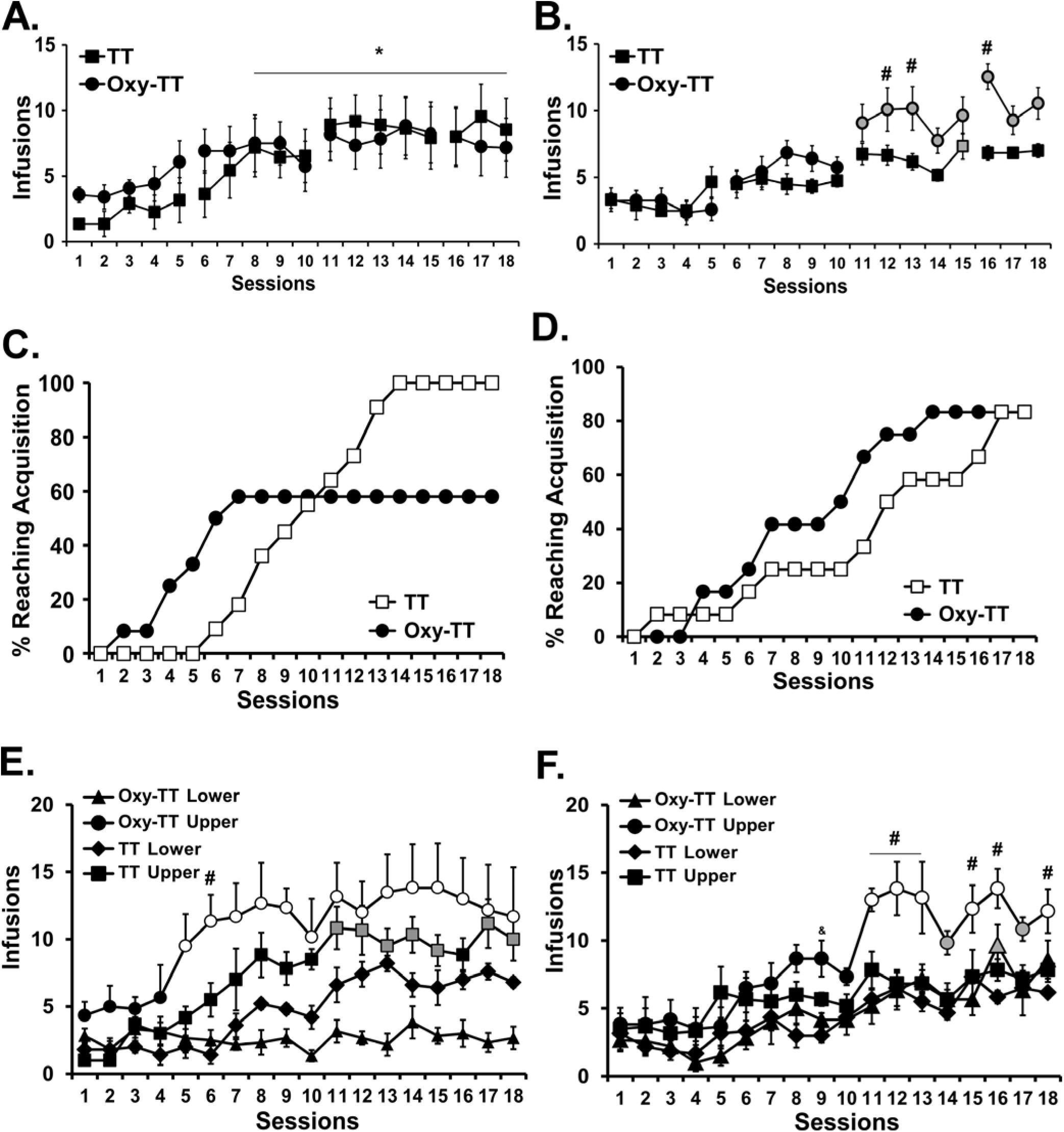
Vaccination attenuates intravenous self-administration of low dose oxycodone. Mean (±SEM) oxycodone infusions obtained by rats in A) Cohort 1 (TT; N=11 and Oxy-TT; N=12) and B) Cohort 2 (TT; N=12 and Oxy-TT; N=11). Post hoc tests: A significant difference from each of the first three sessions, across Groups is indicated with *. Cumulative percentage of each group reaching acquisition criterion (average of 7 infusions across two sessions) in C) Cohort 1 and in D) Cohort 2. Mean (±SEM) infusions obtained by the Upper and Lower (TT Lower N=5) halves of the vaccine groups in E) Cohort 1 and F) Cohort 2. Broken lines between sessions 5 & 6, 10 & 11 and 15 & 16 represent the 70 h weekend interval. Post hoc tests: In panels E and F, the grey symbols indicate a significant difference from the first three sessions within-group. Open symbols reflect significant differences from the first three sessions and between Upper and Lower halves within each vaccine group. A significant difference between vaccine Groups or between the Upper halves of each vaccine group is indicated with #.

### 3.3 Oxy-TT Vaccine Decreases Motivation for Oxycodone Self-Administration

By increasing the workload required for reinforcement, the first PR probe (See details of PR schedule and *j* values in Materials and Methods) in Cohort 1 produced a significant reduction in the number of infusions earned for the Oxy-TT rats only (**Figure 3A**); there was a significant effect of schedule condition [F(1,21)=9.35; *p*=0.006] but not of vaccine Group. The analysis of the saline substitution (**Figure 3B**) confirmed significant effects of vaccine Group [F(1,20)=4.46; *p*=0.0475] and of Drug treatment condition [F(1,20)=28.63; *p*<0.0001], but not of the interaction of factors. Following further training under increasing FR schedule, the TT group obtained more oxycodone infusions under PR (Sessions 64–70) compared with the final FR10 sessions (Sessions 45–63), whereas the Oxy-TT group only increased intake following transition back to the FR1 (Sessions 71–73) condition (**Figure 3C**). The rmANOVA confirmed significant effects of Session [F(8,136)=13.01; *p*<0.0001] and of the interaction of vaccine Group and Session factors [F(8,136)=2.610; *p*=0.0109].

Oxy-TT rats trained under a higher per infusion dose (Cohort 2) also self-administered less oxycodone under the higher workload conditions of a PR schedule (**Figure 3D**). The analysis confirmed an interaction of Group with reinforcement schedule [F(1,20)=19.68; *p*<0.0005]. In the subsequent FR, PR^J2^, PR^J3^, FR transitions fewer infusions were obtained in both PR^J2^ and PR^J3^ relative to each of the FR conditions within the Oxy-TT group (**Figure 3E**). The analysis confirmed a main effect of schedule [F(3,54)=10.45; *p*<0.0001] and interaction of Group with reinforcement contingency [F(3,54)=6.23; *p*=0.001]. Finally, the analysis of the final FR (0.15 mg/kg/inf), FR (0.06 mg/kg/inf), PR^J3^ (0.06 mg/kg/inf) conditions (**Figure 3F**) confirmed a significant impact of the dose/schedule factor [F(2,36)=62.15; *p*<0.0001] and an interaction of this factor with Group [F(2,36)=3.51; *p*=0.0406].

**Figure 3.**
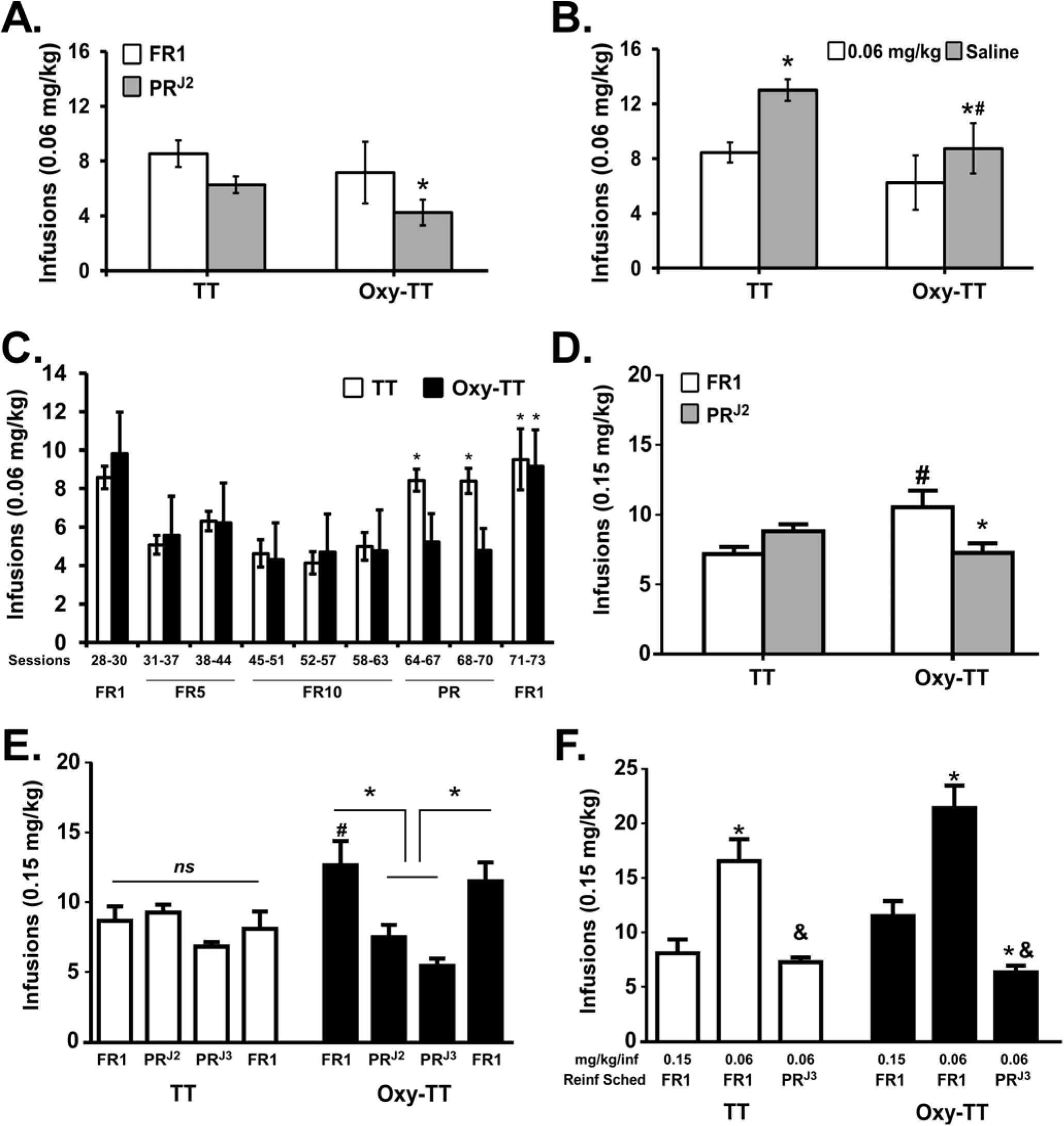
Vaccination decreases motivation for oxycodone self-administration under increased workload. Mean (±SEM) oxycodone infusions obtained by TT (N=11) and Oxy-TT (N=11–12) vaccinated groups in Cohort 1 under A) PR conditions or B) during a saline challenge. Post hoc tests: A significant difference within-group is indicated with * and between the groups with #. C) Training under FR1 to FR10 followed by PR challenges showed that TT (N=11) rats obtained more infusions compared to Oxy-TT (N=8) rats. Post hoc tests: A significant difference from the FR 10 (Sessions 45–63) condition is indicated with *. Mean (±SEM) infusions of oxycodone for Oxy-TT (N= 9–11) and TT (N= 11) vaccinated rats under D) PR conditions and B-C) subsequent challenges with intermittent changes in workload (PR^J2^-PR^J3^). Post hoc tests: A significant difference between vaccine groups for a given schedule of reinforcement condition is indicated by #, a difference from FR (0.15 mg/kg/inf) within group by * and from FR (0.06 mg/kg/inf) within group by &. ns indicates no significant difference was confirmed.

### 3.4 Vaccine Attenuates Antinociceptive Effects of Oxycodone

The effects of oxycodone on nociception were first determined prior to self-administration training in Cohort 1 (N=11–12) and Cohort 2 (N=12) rats. For Cohort 1 (**Figure 4A**), the rmANOVA confirmed significant main effects of vaccine Group [F(1,21)=19.45; *p*=0.0002], of Time [F(4,84)=116; *p*<0.0001], and of the interaction [F(4,84)=8.802; *p*<0.0001]. For Cohort 2 (**Figure 4B**), the analysis also confirmed a significant effect of vaccine Group [F(1,22)=6.36; *p*=0.0194], of Time [F(8,186)=17.1; *p*<0.0001] and of the interaction [F(8,176)=5.39; *p*<0.0001]. The dose-dependent attenuation of the antinociceptive effects of oxycodone in Oxy-TT rats persisted throughout the study (Post-acquisition; **Figure 4A,B**). For Cohort 1 (Week 10), the analysis confirmed a significant effect of Time [F(4,84)=67.26,*p*<0.0001], of Vaccine group [F(1,21)=11.71; *p*=0.0026, and of the interaction [F(4,84)=3.024; *p*=0.0221]. Post hoc analysis confirmed increased tail withdrawal latency in TT rats compared to Oxy-TT during the 30 and 120 min time points. For Cohort 2 (Week 9), the analysis confirmed a significant effect of Time [F(4,88)=33.22; *p*<0.0001], of Vaccine group [F(1,22)=17.6; *p*<0.0005],and of the interaction [F(4,88)=6.499; *p*=0.0001]. Post hoc analysis confirmed tail withdrawal latency significantly increased during the 150 min time point in TT control rats following an injection of 2.0 mg/kg oxycodone. Ultimately, the attenuation of oxycodone antinociception was observed up to 8–10 weeks (data not shown) after the most recent vaccination boost. Finally, Oxy-TT vaccination did not protect against heroin (1.0 mg/kg, s.c.)-induced antinociception which was assessed in Cohort 2 (**Figure 4C**). The rmANOVA confirmed significant effects of vaccine Group [F(1,22)=6.33; *p*<0.0197] and of Time [F(4,88)=28.45; *p*<0.0001].

**Figure 4.**
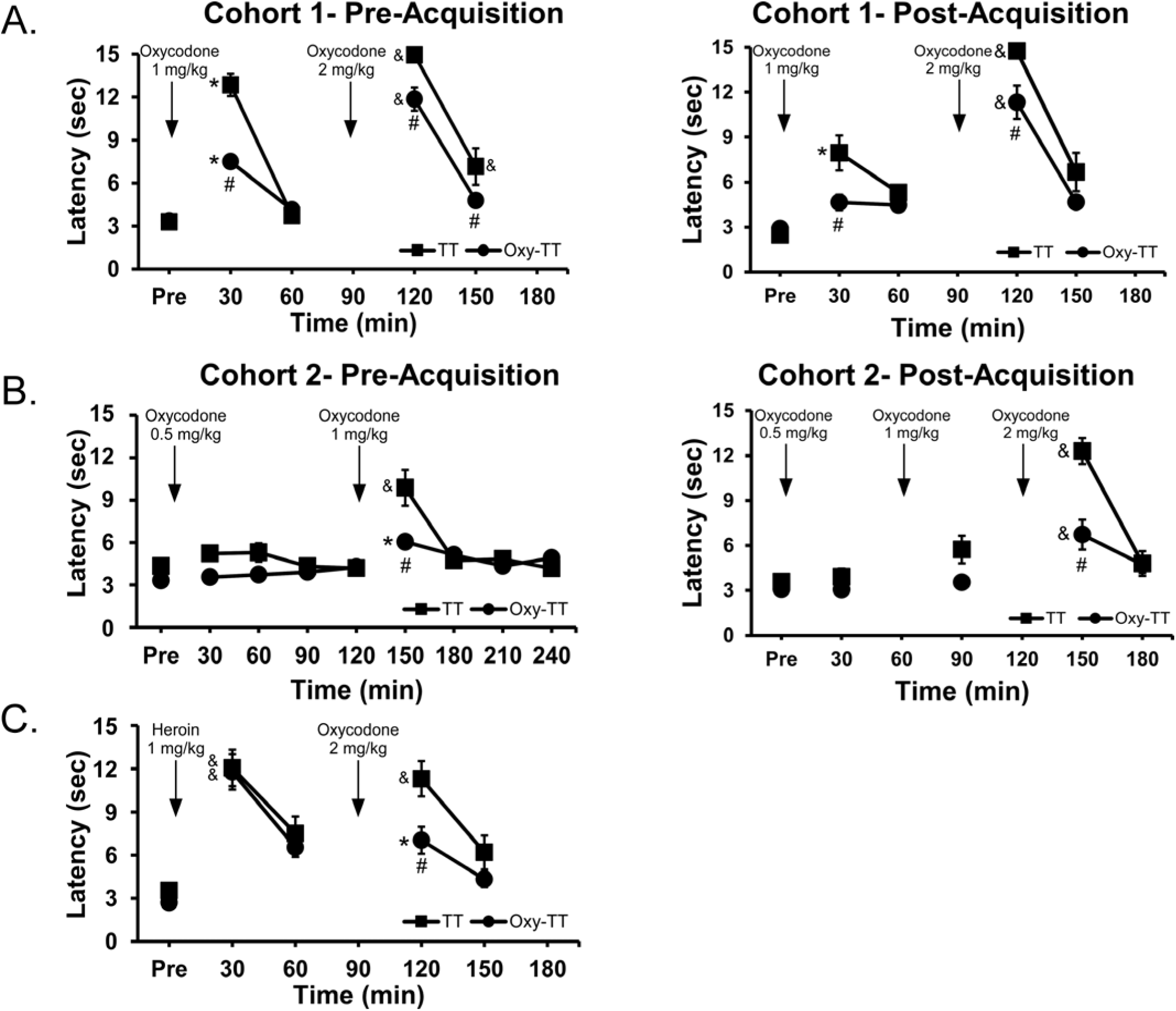
Vaccination ameliorates antinociceptive effect of oxycodone. Mean (±SEM) tail-withdrawal latency in Oxy-TT (N=12) vaccinated rats as compared to TT (N=11–12) control rats prior to and post-IVSA training in A) Cohort 1 and B) Cohort 2. C) In Cohort 1, a non-contingent injection heroin (1 mg/kg, s.c.) showed no difference between groups, but persistence of the difference after oxycodone injection (2 mg/kg, s.c.). Post hoc tests: Significant differences between the groups is indicated with #. Significant differences within group compared with Pre are indicated by * and compared with the most prior observation before a drug injection (60 or 120 min) with &.

### 3.5 Vaccine Decreases Brain Penetration of Oxycodone

Plasma oxycodone (ng/mL) was significantly higher in the Cohort 1 Oxy-TT rats compared to the corresponding TT group following injection of oxycodone (1.0 mg/kg, s.c., **Figure 5A**), which was confirmed by an unpaired t-test [t(20)=7.708, *p*<0.0001]. In Cohort 2, plasma oxycodone was also higher in Oxy-TT rats compared to TT rats (**Figure 5B**) as confirmed by significant effects of vaccination Group, [F(1,22)=73.35; *p*<0.0001]), of Drug condition [F(2,44)=24.13; *p*<0.0001], and of the interaction [F(2.44)=14.98; *p*<0.0001]. Plasma levels of oxycodone were 17- and 25-fold greater in Oxy-TT versus TT at 15 and 30 minutes, respectively, following 1 mg/kg, s.c.. Plasma oxycodone levels in TT rats 30 minutes after injection of 2 mg/kg oxycodone were approximately doubled compared to plasma levels after injection of 1.0 mg/kg oxycodone (**Figure 5B inset**), whereas plasma oxycodone levels in Oxy-TT rats were similar across doses, possibly reflecting the binding capacity of the Abs. Analysis of brain tissue from Cohort 2 after injection of 1.0 mg/kg, s.c., showed significantly lower brain concentrations of oxycodone in Oxy-TT rats compared to TT rats (Figure 5C; [t(19)=4.569, *p*<0.001]). The high concentration of oxycodone in the plasma of Oxy-TT rats is consistent with an interpretation that anti-Oxy Abs sequester free drug in the periphery, preventing or substantially reducing access of the drug to the brain. The oxycodone levels in the TT group decreased by 2-fold from 15 to 30 minutes (after 1.0 mg/kg, s.c.), whereas the decrease of oxycodone levels in the Oxy-TT group was a lower percentage. The ANOVA confirmed effects of Group [F(1,22)=73.35; *p*<0.0001], of Drug Condition [F(2,44)=24.13; *p*<0.0001] and of the interaction of group with dose/time [F(2,44)=14.98; *p*<0.0001]. The Tukey post hoc test confirmed that Oxy-TT and TT oxycodone concentration levels found in plasma under all dose and time post-injection conditions were significantly different.

**Figure 5.**
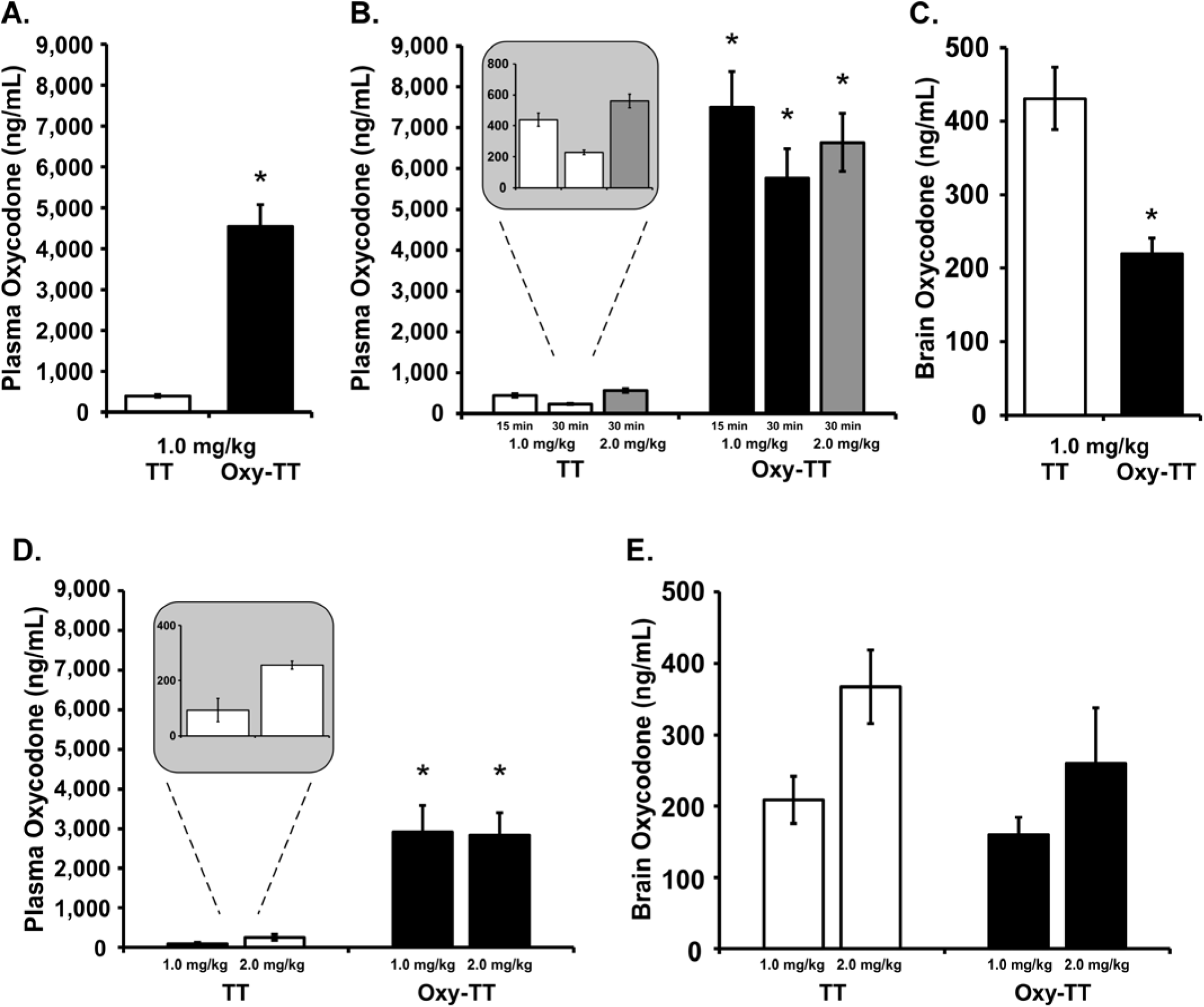
Analysis of oxycodone biodistribution in plasma and brain. A) Mean (±SEM) oxycodone concentrations in plasma, 30 min after injection of oxycodone (1.0 mg/kg, s.c.,) in TT and Oxy-TT groups (N=11 per group) of Cohort 1. B) Mean (±SEM) oxycodone concentrations in plasma at 15 or 30 min after injection of oxycodone (1.0 or 2.0 mg/kg, s.c.) in TT (N=10) and Oxy=TT (N=11; blood could not be obtained from one animal in this experiment) groups of Cohort 2. The insert shows that TT plasma levels at 30 min were twice as high after 2 mg/kg injected versus 1 mg/kg. C) Mean (±SEM) oxycodone concentrations in whole brain tissue at 30 min after injection of oxycodone (1.0 mg/kg, s.c.) in TT (N=10) and Oxy-TT (N=11) groups of Cohort 2. At the end of study in Cohort 1 (Week 29), D) plasma oxycodone levels were significantly higher in OxyTT rats compared to TT (N=5–6) rats 30 min after an injection of oxycodone (1.0–2.0 mg/kg, s.c.), whereas E) brain oxycodone levels showed no difference between OxyTT and TT rats. Post hoc tests: A significant difference between vaccine Groups is indicated with *.

At the end of study in Cohort 1 (Week 29), plasma oxycodone levels were significantly higher in OxyTT (N=5 per dose) rats compared to TT (N=5–6) rats 30 min after an injection of oxycodone (1.0–2.0 mg/kg, s.c.) (**Figure 5D**). The ANOVA confirmed a significant main effect of Vaccine group [F(1,17)=42.69; *p*<0.0001]. Plasma oxycodone levels in TT rats following injection of 2.0 mg/kg were approximately double plasma levels of 1.0 mg/kg oxycodone but again no dose-related differences were observed in the Oxy-TT group. Brain oxycodone levels showed no significant difference between Oxy-TT and TT rats (**Figure 5E**).

## 4. Discussion

This study provides evidence that active vaccination with Oxy-TT provides functional protection against the reinforcing effects of oxycodone. The study demonstrated antinociceptive and pharmacokinetic efficacy following non-contingent drug exposure, thus confirming basic biological activity. The Oxy-TT vaccination resulted in about half of animals failing to acquire stable levels of intravenous self-administration (IVSA) using threshold training conditions and resulted in greater susceptibility of those animals that had established consistent self-administration (under either lower or higher per-infusion dose training conditions) to reduce their intake under a PR schedule of reinforcement. This latter finding, combined with evidence of reduced brain penetration of oxycodone, suggests antidrug vaccination may be effective in synergy with other interventions which help to reduce drug use.

The tail-withdrawal test demonstrated that the Oxy-TT groups were less sensitive to the antinociceptive effect of injected oxycodone in a dose-dependent manner. This was observed prior to the start of self-administration training and throughout the course of the study and included a dose (2.0 mg/kg) that produced a maximal possible effect on the assay in the control group when assessed prior to the start of self-administration (Cohort 1, TT). The post-acquisition test in both Cohorts (**Figure 4A, B**) indicated minor tolerance to the antinociceptive effects of oxycodone (i.e. after the 1.0 mg/kg dose) compared with the pre-acquisition test, most parsimoniously attributed to the oxycodone exposure during self-administration. Importantly, the tail-withdrawal test in Cohort 2 (Week 11) found that while Oxy-TT animals were less sensitive to the effects of *oxycodone*, all animals were equally affected by *heroin* (**Figure 4C**). This shows the selectivity of the protection afforded by anti-oxycodone vaccination and demonstrates that the higher cumulative number of infusions obtained by the Cohort 2 Oxy-TT group did not produce cross-tolerance to heroin, and by inference, differential tolerance to oxycodone. This provides further evidence that the higher IVSA in the Cohort 2 Oxy-TT group did not result in higher ongoing drug exposure to the central nervous system. Similar dose-relationships were observed in the pre-acquisition tail-withdrawal latencies across Cohorts despite different group mean antibody titer. These non-contingent studies therefore highlight the lack of simple correlation of titer with *in vivo* outcome, and confirm antibody hapten binding (as detected via ELISA) does not necessarily equate to functional protection *in vivo*. This is also consistent with a lack of differential protection associated with titer within-Cohort (**Figure 1D**), i.e., associated with the median-split in the distribution of self-administration preference. Analyses of the relationship between antibody titer and behavioral measures (antinociception and self-administration; **Figure S10, S11**) and pharmacokinetics (plasma oxycodone; **Figure S12**) showed a lack of correlation. As a minor caveat, higher binding efficiency in Cohort 1 may have partially compensated for lower titer (**Figure 1C,D**).

The pharmacokinetic experiments found that the Oxy-TT vaccinated rats from each Cohort had approximately 12–15-fold higher plasma oxycodone 30 minutes after 1 mg/kg oxycodone, s.c., and in Cohort 2 this coincided with about half (49%) the brain levels of oxycodone, compared with TT controls, at the end of the study (**Figure 5**). The number of infusions obtained at the end of the acquisition interval in the Cohort 2 study was only 45% higher in the Oxy-TT animals suggesting there may have been a ~26% lower brain oxycodone level in the Oxy-TT group when 0.15 mg/kg/infusion was available under FR1 at the end of the acquisition phase and 34% lower brain oxycodone when 0.06 mg/kg/infusion was available under FR1 near the end of the study. Interestingly, plasma oxycodone levels were dose-dependent in the TT controls (both Cohorts) but not in the Oxy-TT rats, indicating that the saturation point of protection was between 1 and 2 mg/kg. Correspondingly antinociceptive effects of oxycodone were observed after 2 mg/kg, but not 1 mg/kg, s.c., in the Oxy-TT groups.

Acquisition of oxycodone IVSA under the lower per-infusion-dose training (Cohort 1) was associated with significant intra-group differences in the Oxy-TT rats (**Figure 2E**). The Lower half of the distribution obtained fewer infusions than the TT group’s Lower half, while the Upper half of the Oxy-TT group self-administered more oxycodone compared with either half of the TT group. This intra-group difference in IVSA was not associated with any difference in the blunting of the antinociceptive effects of oxycodone either before or after the acquisition training (not shown), nor any significant difference in antibody titer (**Figure 1D**). This lack of difference is consistent with prior studies that have not found strong correlations between individuals’ titer and behavioral responses to drug (LeSage *et al*, 2006). Interestingly, unlike the intra-group differences of oxycodone IVSA there were no significant differences in nociception or the pharmacokinetic measures between Upper and Lower subgroups from either Cohort. This further highlights the importance of investigating vaccine effects on non-lethal oxycodone-related behavior including self-administration.

Another major outcome of this study was the Oxy-TT vaccinated rats that did acquire stable IVSA exhibited greater sensitivity to reduce their drug intake under an increased workload. This was observed in Cohort 2 despite the fact that more infusions were obtained by Oxy-TT rats when only one response was required. When the difficulty was increased by introducing a PR schedule, Oxy-TT rats obtained fewer infusions compared with the TT group and significantly fewer infusions than their own intake under FR1 (**Figure 3E**). In comparison, the TT group did not alter their intake in response to the increased workload. A similar sensitivity of the Oxy-TT group to increased workload was also observed in Cohort 1. Both groups self-administered less oxycodone, compared with the FR1 condition, when infusions were contingent on a higher, fixed number of responses (FR5, then FR10); **Figure 3C**. However, once a PR schedule was introduced the TT group obtained *more* drug because the cumulative response requirement was *lower* than in the prior FR10 sessions (the PR^J2^ required only 9 responses for the fourth infusion, and fewer for the prior three infusions), but the Oxy-TT group did not similarly increase their intake. Upon restoration of the FR1 schedule, the Oxy-TT animals increased to match the TT group, which did not further elevate their intake compared to the intake under PR. These results are both consistent with the Oxy-TT animals being much more sensitive to increasing the demand of obtaining drug infusions. A similar reduction in self-administration under a PR contingency with a cocaine vaccine has been previously reported (Wee *et al*, 2012). This result is behavioral confirmation that the Oxy-TT group experienced less brain exposure during the course of the IVSA training, consistent with a less-dependent state despite the greater number of infusions self-administered. Again, assuming the Oxy-TT rats’ brain oxycodone levels were 49% lower than controls’ (**Figure 5C**), functional exposure under PR was 42% (first PR), 41% (PR^J2^; 0.15 mg/kg/inf and PR^J3^; 0.15 mg/kg/inf) and 44% (PR^J3^; 0.06 mg/kg/inf) lower in the Oxy-TT group. The estimate based on the terminal pharmacokinetic study is, of course, less precise than an assessment of concurrent brain accumulation of drug in real time. However it is, if anything, likely a conservative estimate on two grounds. First, antibody efficacy in the first minutes of exposure under incremental oxycodone infusion conditions, critical to the reinforcing event, is likely higher compared with a single large bolus injection. Second, the antibody titers were likely less than maximum at the end of the study. Overall this suggests that any behavioral compensation for the reduced brain penetration of drug was incomplete and insufficient to produce brain oxycodone levels as high as those achieved by the TT control group.

The results of this study have important implications for the interpretation of the efficacy of antidrug vaccination in pre-clinical models and the application to eventual clinical trials. The results from Cohort 1 illustrate the likely success of prophylactic vaccination prior to any drug experience. That is, a subset of vaccinated animals may not ever acquire high levels of self-administration, particularly given a threshold level of drug exposure. This is consistent with the prior finding of Pravetoni (Pravetoni *et al*, 2014) which used the same 0.06 mg/kg/infusion training dose, although that study also used an increasing Fixed-Ratio across acquisition sessions which complicates interpretation. Delayed acquisition of stable IVSA was also reported in a study of the efficacy of an anti-methamphetamine vaccine (Miller *et al*, 2015). It is also notable that the amount of oxycodone self-administered depended on how much effort was required to obtain each infusion, most pointedly in Cohort 2. Increasing the workload unmasked the fact that the Oxy-TT rats were less motivated to seek oxycodone even if they obtained more infusions under easy access (FR1) conditions. This observation is also consistent with a protective effect of the vaccination, even for individuals who are ingesting psychoactive doses regularly. This makes it more likely that prior results for oxycodone (Pravetoni *et al*, 2014) and methamphetamine (Duryee *et al*, 2009) self-administration was due to the workload increasing across training sessions rather than behavioral extinction.

One caveat to the translational application of this study lies in the focus on non-medical use of a drug that is approved for medical use. Use of such a vaccine as either prophylaxis or as post-cessation therapy in those diagnosed with substance abuse or dependence would have to include close consideration of the diminishment of clinical efficacy, as modeled here with the anti-nociception assay. Such consideration might include the likelihood of the individual requiring opioid medications in the near future and the availability of alternatives, as well as the relative risks of potentially using a higher dose of, e.g., oxycodone, in a very short course of pain management. This highlights the advantages conferred by the specificity of this vaccine, as demonstrated in this study with the antinociceptive efficacy of heroin. Conversely it also underlines the potential need for multiple vaccines to address differing risk profiles with respect to abuse of multiple opioids, e.g. as modelled by Hwang and colleagues (Hwang *et al*, 2018b). For example, high school seniors in the US are 16 times likelier to use any prescription opioid (11 times more likely to use Oxycontin^®^ specifically) than heroin in the past year (Miech *et al*, 2017).

This ratio is cut in half for seniors endorsing past month use, suggesting a change in relative risk from prescription to illicit opioids in the more-frequently using population which would potentially pivot health care priorities from single drug prophylaxis to multi-drug prevention/therapy depending on exposure history.

These results suggest a critical reconsideration of the design and interpretation of human clinical trials in at least two particulars. First, these data encourage consideration of trials for prophylactic efficacy since the Lower half of the Cohort 1 Oxy-TT distribution did not establish a consistent drug-taking pattern. Human drug dependence is a minority outcome among those that sample a given drug (Schramm-Sapyta *et al*, 2009) and it is intuitively obvious that preventing the establishment of dependence from the outset is a sure way to reduce drug-related harms. Second, even though the Upper half of the Oxy-TT distribution in Cohort 2 reached stable, higher patterns of self-administration during acquisition, their intake was more easily reduced compared with the TT animals. The increased intake of oxycodone in the Upper halves of each of the Oxy-TT vaccinated groups might be misinterpreted in a human clinical context as a significant failure, since drug-taking is usually the main outcome measure of clinical trials (Hoogsteder *et al*, 2014; Martell *et al*, 2009). If, as in these preclinical studies, the human users’ drug intake fails to fully compensate for the reduced brain levels of drug this could actually be a treatment success that results in lower dependence over time and possibly in a decreased risk of relapse. This might also facilitate the effects of other therapeutic interventions, similar to the effects of increasing the workload under PR conditions in the present results. In summary, this study showed *in vivo* efficacy of a vaccine directed against oxycodone across pharmacokinetic, non-contingent administration and self-administration assays. The manner by which self-administration was affected provides new insights into how anti-drug vaccines might be tested and applied in the human clinical context.

## Acknowledgements

This study was funded by grants from the USPHS R01 DA035281 (M.A.T.), R01 DA024705 (M.A.T.), UH3 DA041146 (K.D.J.) and F32 AI126628 (C.S.H.), which had no further input on the conduct of the work or the decision to publish results. J.D.N, C.S.H., K.D.J. and M.A.T. designed research; J.D.N, C.S.H. and Y.G. performed research; C.S.H. and K.D.J. contributed new reagents; J.D.N, C.S.H., K.D.J. and M.A.T. analyzed data and wrote the paper.

## References

Anthony JC, Warner LA, Kessler RC (1994). Comparative epidemiology of dependence on tobacco, alcohol, controlled substances and inhalants: Basic findings from the national comorbidity survey. Exp Clin Pyschopharm 2(3): 244–268.

Anton B, Leff P (2006). A novel bivalent morphine/heroin vaccine that prevents relapse to heroin addiction in rodents. Vaccine 24(16): 3232–3240.

Bremer PT, Janda KD (2017). Conjugate Vaccine Immunotherapy for Substance Use Disorder. Pharmacol Rev 69(3): 298–315.

Bremer PT, Schlosburg JE, Lively JM, Janda KD (2014). Injection route and TLR9 agonist addition significantly impact heroin vaccine efficacy. Mol Pharm 11(3): 1075–1080.

Carrera MR, Ashley JA, Zhou B, Wirsching P, Koob GF, Janda KD (2000). Cocaine vaccines: antibody protection against relapse in a rat model. Proc Natl Acad Sci U S A 97(11): 6202–6206.

Cornuz J, Zwahlen S, Jungi WF, Osterwalder J, Klingler K, van Melle G, et al (2008). A vaccine against nicotine for smoking cessation: a randomized controlled trial. PLoS One 3(6): e2547.

Dertadian GC, Maher L (2014). From oxycodone to heroin: two cases of transitioning opioid use in young Australians. Drug Alcohol Rev 33(1): 102–104.

Duryee MJ, Bevins RA, Reichel CM, Murray JE, Dong Y, Thiele GM, et al (2009). Immune responses to methamphetamine by active immunization with peptide-based, molecular adjuvant-containing vaccines. Vaccine 27(22): 2981–2988.

Enga RM, Jackson A, Damaj MI, Beardsley PM (2016). Oxycodone physical dependence and its oral self-administration in C57BL/6J mice. Eur J Pharmacol 789: 75–80.

Evans SM, Foltin RW, Hicks MJ, Rosenberg JB, De BP, Janda KD, et al (2016). Efficacy of an adenovirus-based anti-cocaine vaccine to reduce cocaine self-administration and reacqusition using a choice procedure in rhesus macaques. Pharmacol Biochem Behav 150–151: 76–86.

Haney M, Gunderson EW, Jiang H, Collins ED, Foltin RW (2010). Cocaine-specific antibodies blunt the subjective effects of smoked cocaine in humans. Biol Psychiatry 67(1): 59–65.

Hatsukami DK, Jorenby DE, Gonzales D, Rigotti NA, Glover ED, Oncken CA, et al (2011). Immunogenicity and smoking-cessation outcomes for a novel nicotine immunotherapeutic. Clinical pharmacology and therapeutics 89(3): 392–399.

Hodos W (1961). Progressive ratio as a measure of reward strength. Science (New York, NY) 134: 943–944.

Hoogsteder PH, Kotz D, van Spiegel PI, Viechtbauer W, van Schayck OC (2014). Efficacy of the nicotine vaccine 3’-AmNic-rEPA (NicVAX) co-administered with varenicline and counselling for smoking cessation: a randomized placebo-controlled trial. Addiction 109(8): 1252–1259.

Hwang CS, Bremer PT, Wenthur CJ, Ho SO, Chiang S, Ellis B, et al (2018a). Enhancing Efficacy and Stability of an Antiheroin Vaccine: Examination of Antinociception, Opioid Binding Profile, and Lethality. Mol Pharm 15(3): 1062–1072.

Hwang CS, Smith LC, Natori Y, Ellis B, Zhou B, Janda KD (2018b). Efficacious Vaccine against Heroin Contaminated with Fentanyl. ACS chemical neuroscience.

Kantak KM, Collins SL, Lipman EG, Bond J, Giovanoni K, Fox BS (2000). Evaluation of anti-cocaine antibodies and a cocaine vaccine in a rat self-administration model. Psychopharmacology (Berl) 148(3): 251–262.

Kimishima A, Umihara H, Mizoguchi A, Yokoshima S, Fukuyama T (2014). Synthesis of (−)-Oxycodone. Organic Letters 16(23): 6244–6247.

Kimishima A, Wenthur CJ, Zhou B, Janda KD (2016). An Advance in Prescription Opioid Vaccines: Overdose Mortality Reduction and Extraordinary Alteration of Drug Half-Life. ACS Chemical Biology.

Kimishima A, Wenthur CJ, Zhou B, Janda KD (2017). An Advance in Prescription Opioid Vaccines: Overdose Mortality Reduction and Extraordinary Alteration of Drug Half-Life. ACS Chem Biol 12(1): 36–40.

Kosten TR, Rosen M, Bond J, Settles M, Roberts JS, Shields J, et al (2002). Human therapeutic cocaine vaccine: safety and immunogenicity. Vaccine 20(7–8): 1196–1204.

Leri F, Burns LH (2005). Ultra-low-dose naltrexone reduces the rewarding potency of oxycodone and relapse vulnerability in rats. Pharmacol Biochem Behav 82(2): 252–262.

LeSage MG, Keyler DE, Hieda Y, Collins G, Burroughs D, Le C, et al (2006). Effects of a nicotine conjugate vaccine on the acquisition and maintenance of nicotine self-administration in rats. Psychopharmacology (Berl) 184(3–4): 409–416.

Lindblom N, de Villiers SH, Kalayanov G, Gordon S, Johansson AM, Svensson TH (2002). Active immunization against nicotine prevents reinstatement of nicotine-seeking behavior in rats. Respiration; international review of thoracic diseases 69(3): 254–260.

Lockner JW, Janda KD (eds) (2013). Chapter 2: Immunopharmacotherapy for Nicotine Addiction. The Royal Society of Chemistry.

Mars SG, Bourgois P, Karandinos G, Montero F, Ciccarone D (2014). “Every ‘never’ I ever said came true”: transitions from opioid pills to heroin injecting. Int J Drug Policy 25(2): 257–266.

Martell BA, Orson FM, Poling J, Mitchell E, Rossen RD, Gardner T, et al (2009). Cocaine vaccine for the treatment of cocaine dependence in methadone-maintained patients: a randomized, double-blind, placebo-controlled efficacy trial. Arch Gen Psychiatry 66(10): 1116–1123.

Mavrikaki M, Pravetoni M, Page S, Potter D, Chartoff E (2017). Oxycodone self-administration in male and female rats. Psychopharmacology (Berl) 234(6): 977–987.

Mayer-Blackwell B, Schlussman SD, Butelman ER, Ho A, Ott J, Kreek MJ, et al (2014). Self administration of oxycodone by adolescent and adult mice affects striatal neurotransmitter receptor gene expression. Neuroscience 258: 280–291.

Miech RA, Johnston LD, O’Malley PM, Bachman JG, Schulenberg JE (2017). Monitoring the Future national survey results on drug use, 1975-2016. Volume I, Secondary school students Institute for Social Research, The University of Michigan: Ann Arbor, MI, p 683.

Miller ML, Aarde SM, Moreno AY, Creehan KM, Janda KD, Taffe MA (2015). Effects of active anti-methamphetamine vaccination on intravenous self-administration in rats. Drug Alcohol Depend 153: 29–36.

Miller ML, Moreno AY, Aarde SM, Creehan KM, Vandewater SA, Vaillancourt BD, et al (2013). A methamphetamine vaccine attenuates methamphetamine-induced disruptions in thermoregulation and activity in rats. Biol Psychiatry 73(8): 721–728.

Moreno AY, Azar MR, Warren NA, Dickerson TJ, Koob GF, Janda KD (2010). A critical evaluation of a nicotine vaccine within a self-administration behavioral model. Mol Pharm 7(2): 431–441.

Nguyen JD, Bremer PT, Ducime A, Creehan KM, Kisby BR, Taffe MA, et al (2016). Active vaccination attenuates the psychostimulant effects of alpha-PVP and MDPV in rats. Neuropharmacology 116: 1–8.

Nguyen JD, Bremer PT, Hwang CS, Vandewater SA, Collins KC, Creehan KM, et al (2017a). Effective active vaccination against methamphetamine in female rats. Drug Alcohol Depend 175: 179–186.

Nguyen JD, Grant Y, Creehan KM, Vandewater SA, Taffe MA (2017b). Escalation of intravenous self-administration of methylone and mephedrone under extended access conditions. Addict Biol 22(5): 1160–1168.

Ohia-Nwoko O, Kosten TA, Haile CN (2016). Animal Models and the Development of Vaccines to Treat Substance Use Disorders. Int Rev Neurobiol 126: 263–291.

Pravetoni M, Le Naour M, Harmon TM, Tucker AM, Portoghese PS, Pentel PR (2012). An oxycodone conjugate vaccine elicits drug-specific antibodies that reduce oxycodone distribution to brain and hotplate analgesia. J Pharmacol Exp Ther 341(1): 225–232.

Pravetoni M, Pentel PR, Potter DN, Chartoff EH, Tally L, LeSage MG (2014). Effects of an oxycodone conjugate vaccine on oxycodone self-administration and oxycodone-induced brain gene expression in rats. PLoS One 9(7): e101807.

Raleigh MD, Laudenbach M, Baruffaldi F, Peterson SJ, Roslawski MJ, Birnbaum AK, et al (2018). Opioid Dose- and Route-Dependent Efficacy of Oxycodone and Heroin Vaccines in Rats. J Pharmacol Exp Ther 365(2): 346–353.

Raleigh MD, Peterson SJ, Laudenbach M, Baruffaldi F, Carroll FI, Comer SD, et al (2017). Safety and efficacy of an oxycodone vaccine: Addressing some of the unique considerations posed by opioid abuse. PLoS One 12(12): e0184876.

Richardson NR, Roberts DC (1996). Progressive ratio schedules in drug self-administration studies in rats: a method to evaluate reinforcing efficacy. Journal of Neuroscience Methods 66(1): 1–11.

Schlosburg JE, Vendruscolo LF, Bremer PT, Lockner JW, Wade CL, Nunes AA, et al (2013). Dynamic vaccine blocks relapse to compulsive intake of heroin. Proc Natl Acad Sci U S A 110(22): 9036–9041.

Schramm-Sapyta NL, Walker QD, Caster JM, Levin ED, Kuhn CM (2009). Are adolescents more vulnerable to drug addiction than adults? Evidence from animal models. Psychopharmacology (Berl) 206(1): 1–21.

Secci ME, Factor JA, Schindler CW, Panlilio LV (2016). Choice between delayed food and immediate oxycodone in rats. Psychopharmacology (Berl) 233(23–24): 3977–3989.

Segal DS, Mandell AJ (1974). Long-term administration of d-amphetamine: progressive augmentation of motor activity and stereotypy. Pharmacol Biochem Behav 2(2): 249–255.

Stowe GN, Vendruscolo LF, Edwards S, Schlosburg JE, Misra KK, Schulteis G, et al (2011). A vaccine strategy that induces protective immunity against heroin. Journal of medicinal chemistry 54(14): 5195–5204.

Taffe MA (2015). Drug abuse scientists should use social media to engage the public because their primary translational product is information. Drug Alcohol Depend 154: 315–319.

Wade CL, Vendruscolo LF, Schlosburg JE, Hernandez DO, Koob GF (2015). Compulsive-like responding for opioid analgesics in rats with extended access. Neuropsychopharmacology 40(2): 421–428.

Wee S, Hicks MJ, De BP, Rosenberg JB, Moreno AY, Kaminsky SM, et al (2012). Novel cocaine vaccine linked to a disrupted adenovirus gene transfer vector blocks cocaine psychostimulant and reinforcing effects. Neuropsychopharmacology 37(5): 1083–1091.

Zhang Y, Brownstein AJ, Buonora M, Niikura K, Ho A, Correa da Rosa J, et al (2015). Self administration of oxycodone alters synaptic plasticity gene expression in the hippocampus differentially in male adolescent and adult mice. Neuroscience 285: 34–46.

Zhang Y, Picetti R, Butelman ER, Schlussman SD, Ho A, Kreek MJ (2009). Behavioral and neurochemical changes induced by oxycodone differ between adolescent and adult mice. Neuropsychopharmacology 34(4): 912–922.

